# Fungal symbionts produce prostaglandin E_2_ to promote their intestinal colonization

**DOI:** 10.1101/477117

**Authors:** Tze Guan Tan, Ying Shiang Lim, Alrina Tan, Royston Leong, Norman Pavelka

## Abstract

*Candida albicans* is a ubiquitous fungal symbiont that resides on diverse human barrier surfaces. Both mammalian and fungal cells can convert arachidonic acid into the lipid mediator, prostaglandin E2 (PGE_2_), but the physiological significance of fungal-derived PGE_2_ remains elusive. Here we report that a *C. albicans* mutant deficient in PGE_2_ production suffered a loss of competitive fitness in the murine gastrointestinal (GI) tract and that PGE_2_ supplementation mitigated this fitness defect. Impaired fungal PGE_2_ production affected neither the *in vitro* fitness of *C. albicans* nor hyphal morphogenesis and virulence in either systemic or mucosal infection models. Fungus-derived PGE_2_ improved intra-GI fitness of *C. albicans* by diminishing the killing of *C. albicans* by phagocytes. Consequently, ablation of colonic phagocytes abrogated the fitness boost conferred by fungal PGE_2_. These observations suggest that *C. albicans* has evolved the capacity to produce PGE_2_ from arachidonic acid, a host-derived precursor, to promote its own colonization of the host gut. Analogous mechanisms might undergird host-microbe interactions of other symbiont fungi.

**Author Summary:** *Candida albicans* is a symbiont fungus that resides in the gut of a majority of people without provoking disease. However, resident *C. albicans* can bloom and turn pathogenic in a subset of individuals who are immunocompromised due to infections or chemotherapy or who suffer a disruption of their intestinal microbial community due to antibiotic use. However, the fungal and host factors that regulate the fitness of *C. albicans* as a symbiont or an invasive pathogen remain poorly understood. Here we focused on the physiological role of fungus-derived prostaglandin E2 (PGE_2_) in the fitness of *C. albicans* using a PGE_2_-deficient *C. albicans* strain and mouse models of infections and intestinal symbiosis. We found that fungal PGE_2_, contrary to previously described functions of promoting virulence, played no role in fungal pathogenicity *in vivo*. Instead, fungal PGE_2_ specifically augmented the ability of *C. albicans* to colonize the gut, in part by reducing fungal killing by intestinal phagocytes. Our results suggest that fungal PGE_2_ synthetic pathways may be prophylactically targeted in individuals susceptible to invasive infections.

## Introduction

*Candida albicans* is one of the most successful fungal symbionts in humans, colonizing 40-80% of individuals in industrialized nations and typically representing the predominant species within the fungal microbiota [1, 2]. In healthy people, *C. albicans* dwells on diverse barrier sites of the body, including the oral cavity, skin, female reproductive tract, and the intestines, where it does not cause symptomatic disease [1, 2]. However, *C. albicans* can turn pathogenic and result in mucosal or life-threatening invasive bloodstream infections under a variety of conditions that compromise host immunity, damage barrier surfaces, or disrupt the microbiota [3, 4]. Molecular profiling studies have identified *C. albicans* strains in the gastrointestinal (GI) tract as the provenance of hematogenous isolates during systemic infection [5, 6]. Understanding the fungal and host factors that modulate the ability of *C. albicans* to thrive in different host niches as a pathogen or symbiont could therefore inform our development of new methods of combatting human fungal infections.

Screens of *C. albicans* mutants in mice have revealed several fungal pathways that mediate colonization of the host gut. Salient pathways include the regulation of the acquisition of iron and nutrients, in particular carbon and nitrogen, cell wall remodeling that augments fungal adhesion to intestinal mucins, and morphogenetic changes [7–14]. Fungal colonization is, in turn, restricted by the host immune system. Intestinal epithelial cells (IECs) produce antimicrobial peptides (AMPs) that kill *C. albicans* [4], and gut-resident CX_3_CR1^+^ macrophages control the growth of *C. albicans* and other symbiont fungi in mice [15]. Accordingly, some of the aforementioned genetic programs in *C. albicans* might also promote intestinal symbiosis by calibrating the host immune response towards the fungus or *vice versa*. For instance, cell wall remodeling can mask the exposure of β-glucan, an essential component of fungal cell walls, to host immunological sensing via dectin-1 [16, 17], and recognition of surface β-glucan by dectin-1 is strongly correlated with the competitive fitness of *Candida* species in the murine intestine [13]. In addition, *C. albicans* upregulates the key morphogenetic regulator, *EFG1*, upon colonizing the gut, and *EFG1* is required for *C. albicans* to overcome host immunological pressures to persist in the GI tract long-term [18].

Prostaglandin E2 (PGE_2_) is a lipid mediator produced by mammalian cells that exerts pleiotropic effects on physiology, in particular the modulation of immunological responses [19]. For instance, PGE_2_ suppresses the maturation of dendritic cells and restrains the phagocytic activity of monocytes, macrophages, and neutrophils [19, 20]. Depending on the tissue context, PGE_2_ can either promote [21–24] or suppress [25–27] pro-inflammatory CD4^+^ T helper (Th)-1 and −17 responses, which help rein in *C. albicans* infections in the bloodstream and at barrier surfaces respectively [28].

Of note, many fungal species, including *C. albicans*, also produce PGE_2_ *in vitro* when supplied exogenously with the host-derived precursor, arachidonic acid (AA) [29]. *C. albicans* and most fungal species lack AA, which is derived from mammalian membrane phospholipids – but *C. albicans* can induce host cells to release AA [30]. Moreover, fungus-derived PGE_2_ exhibits immunomodulatory properties analogous to those of mammalian PGE_2_ *in vitro* [31] and *in vivo* [32]. PGE_2_ has also been reported to promote yeast-hyphae morphological switching [31, 33] and biofilm formation *in vitro* [34] – traits commonly associated with fungal virulence. Given that the fungus is more typically associated with the host as a benign symbiont and inter-individual transmission often occurs in the absence of overt infection [6, 35], it seems unlikely, however, that *C. albicans* has evolved the cellular machinery for synthesizing PGE_2_ from an exogenous substrate for the sole purpose to increase its pathogenic potential or to cause immune-related disorders in the host. We thus posit that fungus-derived PGE_2_ might mediate asymptomatic colonization by *C. albicans* of the mammalian intestinal environment. Indeed, we found that a *C. albicans* mutant with impaired PGE_2_ production, while not exhibiting any defects in *in vitro* fitness or *in vivo* virulence, suffered a loss of competitive fitness relative to the wildtype (WT) strain in the murine intestine. We further report that fungal PGE_2_ promotes intestinal symbiosis by *C. albicans* in part by reducing its killing by phagocytes.

## Results

### PGE_2_-defective *C. albicans* mutant exhibits no fitness defects *in vitro*

PGE_2_ production has been described for *C. albicans* and other pathogenic fungi such as *Cryptococcus neoformans* and *Aspergillus* species [29] but not for symbiont fungi or other *Candida* species, many of which, like *C. albicans*, can be found in the gut as well. We thus measured *in vitro* PGE_2_ production across an array of symbiont fungi, including various *Candida* species and *Saccharomyces cerevisiae*, as well as a fungus not typically associated with a host, *Schizosaccharomyces pombe*. Fungal PGE_2_ production varied across species, but all species tested, secreted PGE_2_ when supplied with exogenous AA (Fig 1A), although *S. pombe* did so to a lower extent, implying a PGE_2_ biosynthetic pathway conserved across the majority of fungal species.

**Fig 1.**
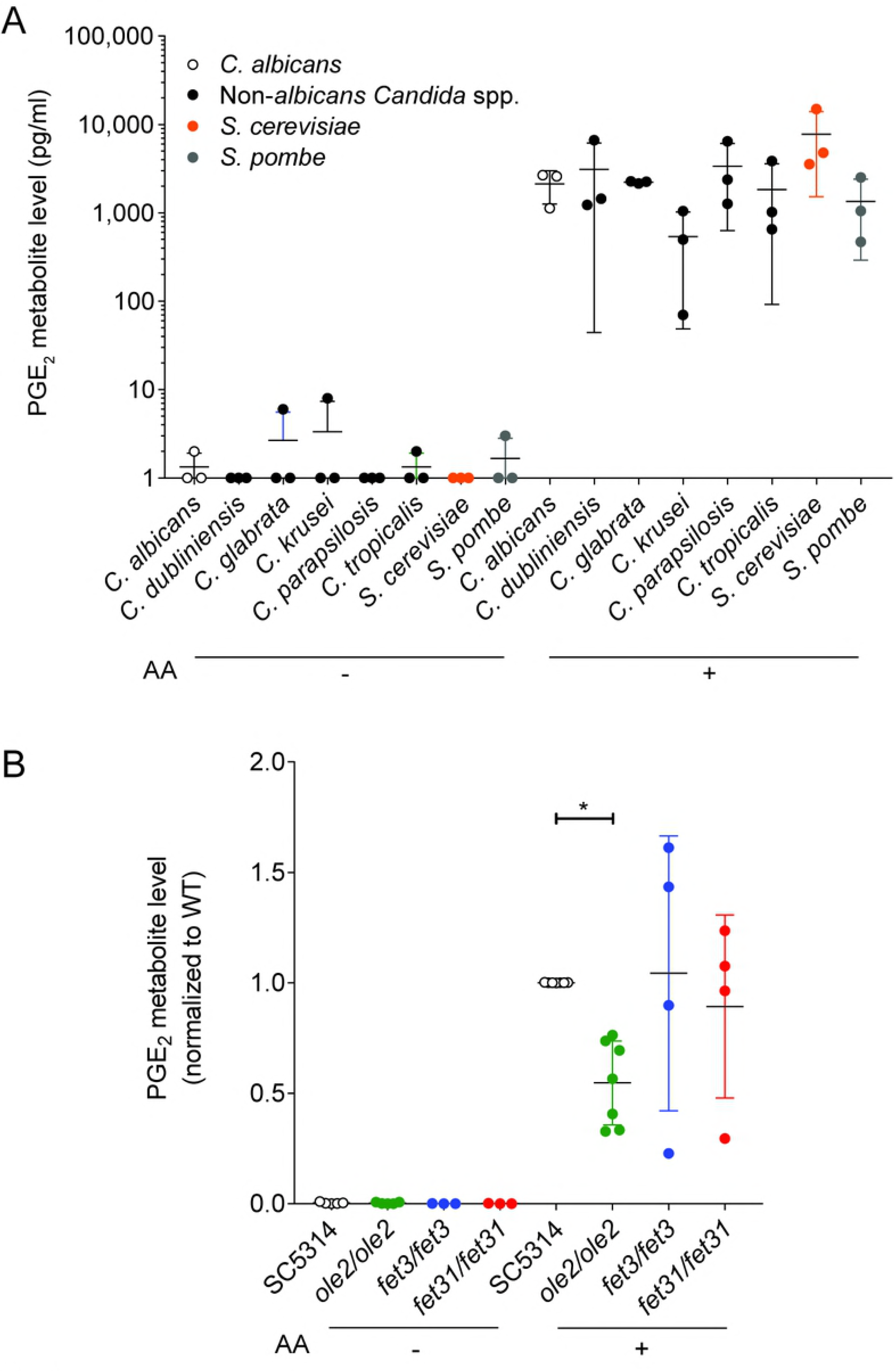
*In vitro* PGE_2_ production by various *C. albicans* strains. Levels of PGE_2_ metabolite, a stable derivative of PGE_2_, in the supernatants of *in vitro* cultures of various fungal species (A) or *C. albicans* strains (B), were assayed by ELISA. Arachidonic acid (AA) was added (+) or not (-) to the cultures. Data pooled from 3-7 independent experiments. Mean ± s.d.. *, *p* < 0.05; One-way ANOVA with Dunnett’s multiple comparisons test.

We next sought a *C. albicans* mutant with altered PGE_2_ production which we could use to study the physiological function of fungal-derived PGE_2_. Although the PGE_2_ biosynthetic pathway in *C. albicans* and other fungi remains largely uncharacterized, three genes, *OLE2*, *FET3*, and *FET31*, have been described to promote PGE_2_ production *in vitro* [36, 37]. We generated knockout (KO) mutants for each of these genes and assessed the mutants’ capacities for PGE_2_ secretion *in vitro* at 37 °C. As expected, PGE_2_ production by all fungal strains was strictly dependent on the provision of exogenous AA (Fig 1B). Because only the *ole2/ole2* mutant exhibited reproducibly diminished PGE_2_ secretion, we focused on this mutant for all subsequent experiments.

To determine whether fungal PGE_2_ modifies fungal fitness *in vitro*, we maintained either the WT or the *ole2/ole2* strain in competition with a fluorescently tagged WT strain (SC5314-dTomato) in serially passaged liquid batch cultures. The abundance of each strain relative to that of SC5314-dTomato was determined at regular intervals by plating the cultures on yeast peptone dextrose (YPD) agar followed by scoring of fluorescent and non-fluorescent colonies. The fitness of each strain can in turn be quantified by deriving a fitness coefficient from the rate of change of the relative abundance of each strain [13]. Even after 80 h (equivalent to ~44 and 63 generations for growth in YNB and YPD, respectively) of serial subculturing, we detected no differences in competitive fitness between the *ole2/ole2* and WT strains in both a nutrient-rich YPD broth and a nutrient-poor yeast nitrogen base (YNB) media (Fig 2). Therefore fungal PGE_2_ does not appear to be important for fitness *in vitro*.

**Fig 2.**
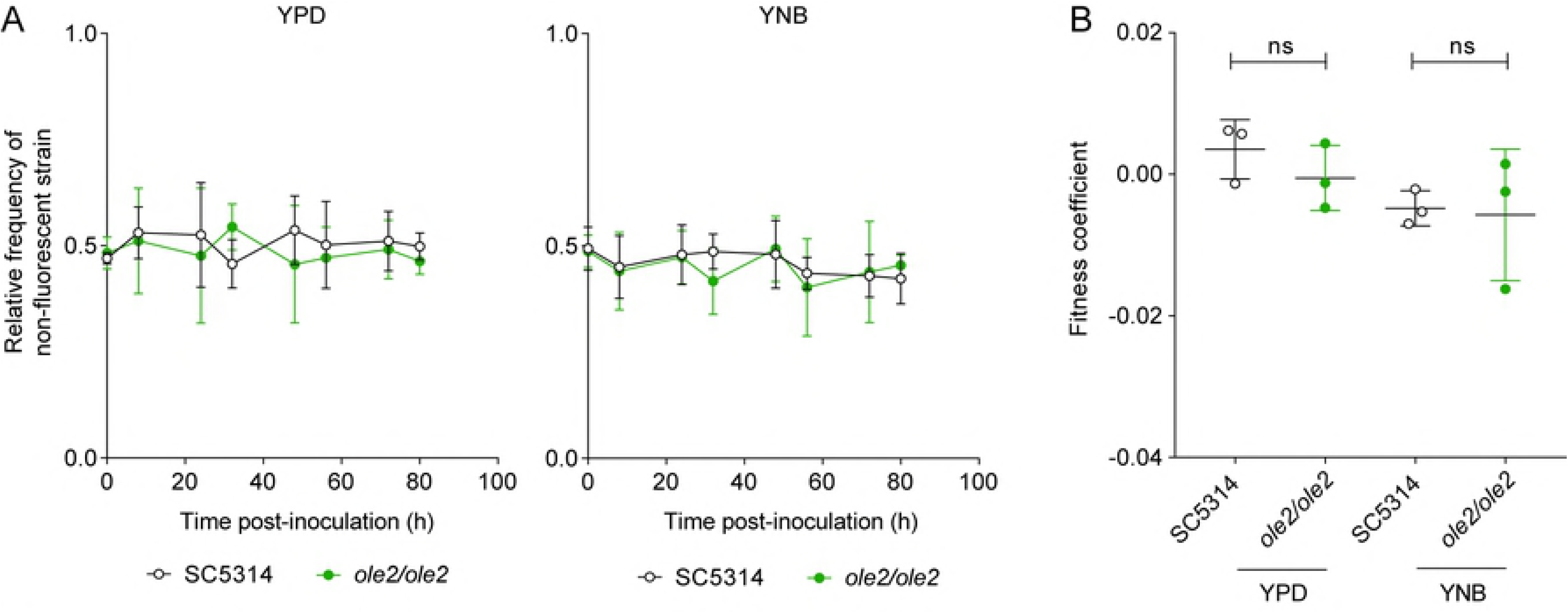
Fungal PGE_2_ is not required for *in vitro* competitive fitness of *C. albicans*. WT *C. albicans* or the *ole2/ole2* mutant was grown in rich (YPD) or minimal (YNB) media in competition with a fluorescently-tagged WT strain at a 1:1 ratio in serial batch cultures for 80 h. (A) Relative frequencies of the WT or *ole2/ole2* strain throughout the serial passages. (B) Fitness coefficients of the WT or *ole2/ole2* strain, determined as outlined in Materials and Methods. Data pooled from 3 independent experiments. Mean ± s.d. (A). ns, not significant. Student’s *t* test.

### Fungal PGE_2_ is not required for virulence

Since PGE_2_ was reported to increase filamentation and biofilm formation *in vitro*, we asked whether defective PGE_2_ synthesis by the *ole2/ole2* mutant influences its virulence. Surprisingly, the *ole2/ole2* mutant was fully capable of hyphal formation *in vitro*, both in response to fetal calf serum (FCS) and when plated on Spider agar (Fig 3A-B). Moreover, the ability of *C. albicans* to damage host cells, evidenced by the release of lactate dehydrogenase from infected murine macrophages, was not significantly altered by fungal PGE_2_ (Fig 3C). Nonetheless, there exist occasional discrepancies between *in vivo* virulence and *in vitro* pathogenic traits [38]. To clarify the role of fungus-derived PGE_2_ during *in vivo* infections, we adopted murine models of hematogenously disseminated candidiasis and mucosal candidiasis of the oropharyngeal cavity and vaginal tract, all of which engage distinct anti-fungal immunological responses [28, 39, 40]. The *ole2/ole2* mutant manifested comparable virulence as the WT strain in all tissues tested, across a variety of pathological parameters, including overall survival, fungal burden, weight loss, and neutrophil recruitment (Fig 3D-F, S1 Fig). Hence these *in vitro* and *in vivo* data are not consistent with a role for fungal PGE_2_ in mediating *C. albicans* virulence.

**Fig 3.**
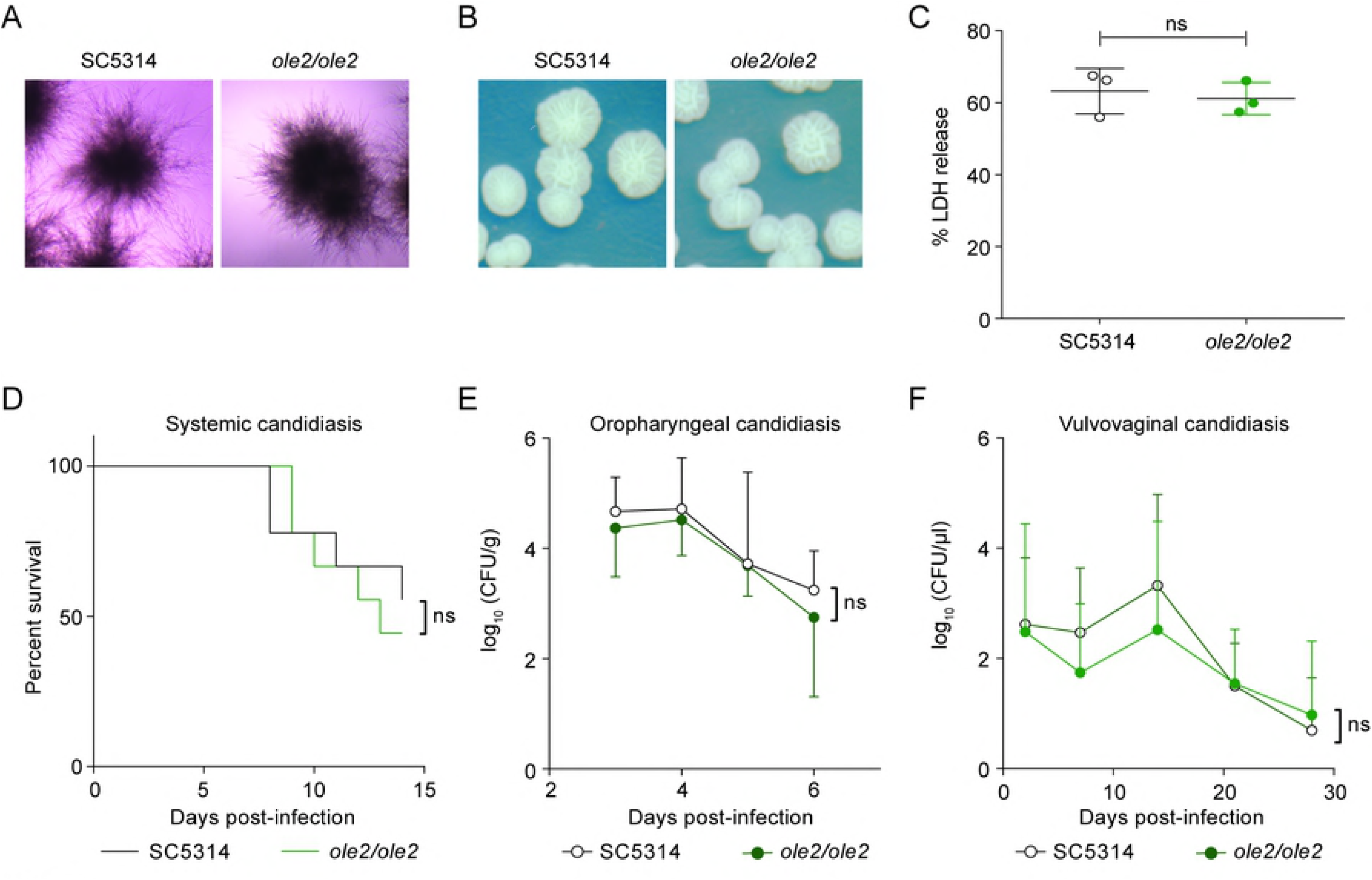
Fungal PGE_2_ is not required for virulence. (A-B) Hyphae formation of the WT and *ole2/ole2 C. albicans* strains in response to fetal calf serum (FCS) (A) or when plated on Spider agar (B). (C) *In vitro* cytotoxicity of the indicated fungal strains to J774A.1 murine macrophages, determined by quantifying the release of LDH from the macrophages. Data presented as a percentage of the maximal amount of LDH released by completely lysed cells (D) Survival of mice infected intravenously with 2 × 10^5^ cells of the indicated *C. albicans* strains. (E-F) Kinetics of the fungal CFUs in the tongue (E) and vaginal lavage fluid (F) of mice infected sublingually (E) or intravaginally (F) with the indicated fungal strains. Data pooled from 2 independent experiments. Mean ± s.d. (C, E-F). *n*=8-10 mice/group. ns, not significant. Student’s *t* test (C, E-F); Log-rank test (D).

### Fungal PGE_2_ promotes intestinal colonization

*C. albicans* is most commonly associated with the host as a symbiont in the gut. We therefore sought to evaluate whether fungal PGE_2_ contributes to the fitness of *C. albicans* in the intestines. Mice are usually refractory to colonization by *C. albicans*, but can be induced to harbor high titers of fungi for sustained durations if treated with antibiotics to deplete their indigenous microbiota [4, 41]. Colonization of the murine GI tract with *C. albicans* represents a *bona fide* model for fungal symbiosis with the mammalian host because it does not lead to symptomatic disease and invasive hyphae are rarely detected within the intestinal fungal population [7, 12, 41]. When the *ole2/ole2* mutant was introduced into mice in competition with SC5314-dTomato, the mutant strain was significantly compromised in its competitive fitness, starting as early as 3 days post-inoculation (Fig 4A-B, S2A Fig). To determine if the loss of competitive fitness of the *ole2/ole2* strain in the gut is PGE_2_-dependent, we supplemented *C. albicans*-colonized mice daily with 16,16-dimethyl-PGE_2_ (dmPGE_2_), a stable analog of PGE_2_. Indeed, dmPGE_2_ treatment partially restored the competitive fitness of the *ole2/ole2* mutant (Fig 4C, S2B-C Fig), in support of a role for fungal PGE_2_ in mediating intestinal fitness of *C. albicans* under steady-state conditions. In contrast, dmPGE_2_ provision did not further enhance the competitive fitness of the WT strain. Taken together, these data indicate that fungal PGE_2_ exerts a specific function for *C. albicans*: promote its colonization of the host GI tract.

**Fig 4.**
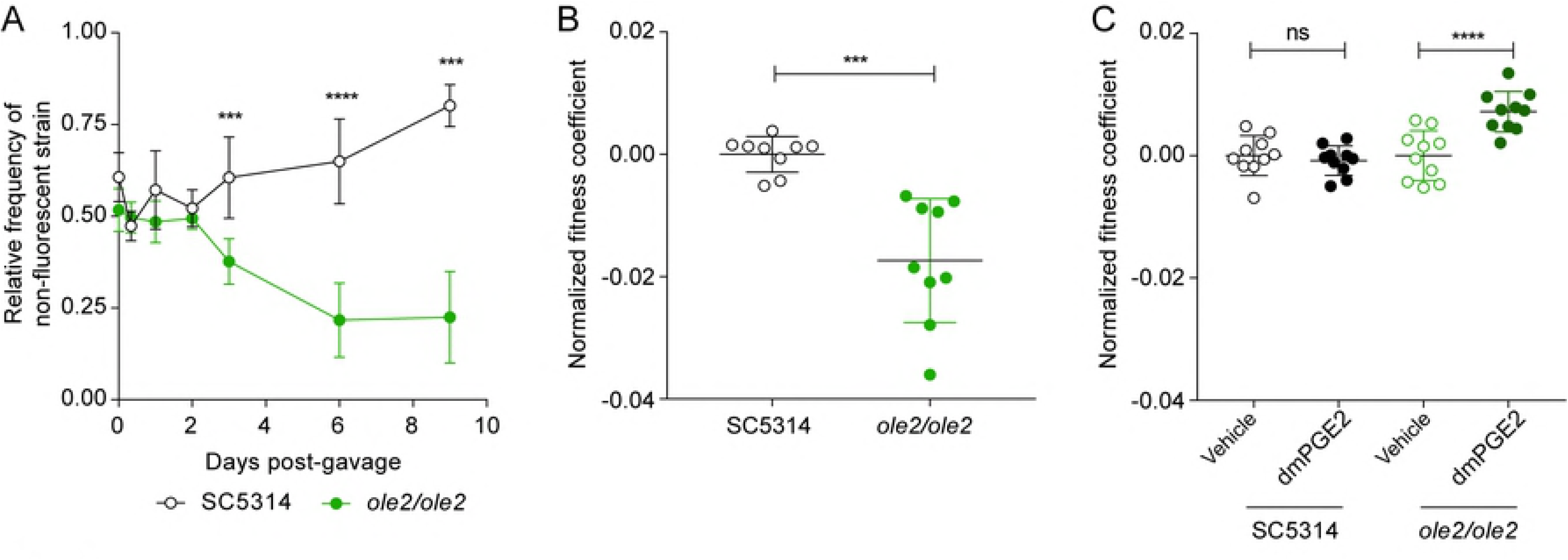
Fungal PGE_2_ promotes intestinal colonization by *C. albicans*. Antibiotics-treated mice were gavaged with the indicated *C. albicans* strains and stools were plated on YPD agar at regular intervals over 9 days. (A-B) Relative frequencies (A) and fitness coefficients, normalized to the mean of the WT coefficients (B), of the indicated fungal strains in the intestines of gavaged mice. (C) Fitness coefficients of the indicated fungal strains in mice supplemented with vehicle or 16,16-dimethyl-PGE_2_ (dmPGE_2_). Coefficients for each fungal strain were normalized to the mean of the respective vehicle coefficients. Fitness coefficients were determined as described in Materials and Methods. Data pooled from 2 independent experiments. Mean ± s.d.. *n*=9-10 mice/group. ****, *p* < 0.0001; ***, *p* < 0.001. Student’s *t* test (A-B); Two-way ANOVA with Sidak’s multiple comparisons test (C).

### Fungal PGE_2_ does not affect AMP production or fungal resistance to AMPs

AMPs such as mouse cathelicidin-related antimicrobial peptide (mCRAMP), encoded by *Camp*, have been demonstrated to mediate intestinal resistance to *C. albicans* colonization [4]. We postulated that fungal PGE_2_ might promote intestinal fitness of *C. albicans* by influencing AMP production from IECs or fungal resistance to AMPs. To address the former hypothesis, we stimulated human colonic IEC lines with WT *C. albicans* or the *ole2/ole2* mutant, but observed no difference in the secretion of LL-37, the human ortholog of mCRAMP, by the IEC lines (S3A Fig). Indeed, infection of the IEC lines by fungi *in vitro* did not induce appreciable production of LL-37 above the background level. We also assessed *Camp* transcript levels in the colons of mice colonized with either WT or *ole2/ole2 C. albicans*, but found no difference in the quantities of *Camp* mRNA induced by the two fungal strains *in vivo* (S3B Fig). Finally, we exposed fungi grown in the presence or absence of PGE_2_ to mCRAMP *in vitro* to evaluate the effects of PGE_2_ on fungal resistance to AMPs. Both the WT and *ole2/ole2* strains manifested comparable susceptibility to mCRAMP, and the addition of PGE_2_ did not significantly alter fungal resistance to mCRAMP (S3C Fig). Hence the collective evidence does not support a role for fungus-derived PGE_2_ in altering AMP production from IECs or fungal resistance to AMPs.

### Fungal PGE_2_ does not modulate β-glucan exposure and dectin-1 signaling

In an earlier report, we had demonstrated that the extent of surface β-glucan exposure by fungal cells and their consequent triggering of dectin-1 signaling correlate strongly with the *in vivo* intestinal fitness of *Candida* species [13]. We thus sought to determine if fungal PGE_2_ modulates fungal fitness in the murine gut by tuning β-glucan exposure on fungal cell wall surfaces. WT *C. albicans* exposed significantly less β-glucan than the *ole2/ole2* strain in the absence of PGE_2_ (S4A Fig). When PGE_2_ production was induced by the addition of exogenous AA, the *ole2/ole2* mutant still exposed more β-glucan on its surface, though the difference in exposure between the two strains was subtle (S4A Fig). To determine if the preceding differences in β-glucan exposure have functional consequences, we measured the ability of the two fungal strains to induce dectin-1 signaling in a hDectin-1b reporter cell line. In contrast to our data on β-glucan surface exposure, the WT and *ole2/ole2* strains elicited comparable levels of dectin-1 signaling, with or without AA (S4B Fig). The discrepancy between β-glucan exposure and dectin-1 signaling could be due to structural differences in surface β-glucan between the two strains that affect recognition and signaling by dectin-1 but are not detectable by flow cytometry. In sum, fungal PGE_2_ does not appear to mediate fungal fitness in the GI tract by modulating the β-glucan-dectin-1 signaling axis.

### Fungus-derived PGE_2_ reduces killing of *C. albicans* by intestinal phagocytes

Since intestinal macrophages limit fungal colonization of the gut [15] and tissue-resident phagocytes are required for control of fungal infection in various tissues [28, 42, 43], we postulated that fungal PGE_2_ might potentiate fungal fitness by acting on the fungi themselves and/or host tissular phagocytes to improve the ability of *C. albicans* to evade killing by phagocytes. To elucidate the relationship between fungal PGE_2_, tissular phagocytes, and fungal fitness in the host, we assessed the competitive fitness of the WT and *ole2/ole2* strains in mice depleted of phagocytes. We first treated mice with clodronate-containing liposomes, which is toxic to a variety of phagocytic leukocytes, including macrophages and dendritic cells (DCs) [44]. Clodronate treatment depleted colonic CD11b^+^ dendritic cells (DCs) but not macrophages, CD11b^−^ DCs, or non-phagocytic leukocytes (S5A-B Fig). The poor efficiency of macrophage ablation in clodronate-treated mice could possibly arise from differential accessibility and susceptibility of macrophages in the colon to intraperitoneally-administered clodronate relative to those from other tissue sites. Nonetheless, clodronate treatment was sufficient to abrogate the fitness defect of the *ole2/ole2* mutant vis-à-vis the WT strain that was observed in control mice, thereby implicating CD11b^+^ DCs as another tissue-resident phagocytic population that restricts *C. albicans* in the gut (Fig 5A-B, S6A Fig). We further validated the preceding results by repeating the fungal competitive colonization experiment in a genetic model of phagocyte depletion. CCR2 is a chemokine receptor required for efficient reconstitution of the intestinal macrophage and CD11b^+^ DC compartments [45, 46]. Consequently, *Ccr2*-KO mice display a paucity of both macrophages and CD11b^+^ DCs but not CD11b^−^ DCs or lymphocytes in the colon (S5C-D Fig). In agreement with our findings in clodronate-treated mice, *Ccr2* deficiency abolished the fitness difference between the WT and *ole2/ole2* strains observed in WT mice (Fig 5C-D, S6B Fig). Our results collectively support a role for fungal PGE_2_ in reducing the eradication of *C. albicans* by host intestinal phagocytes, including macrophages and CD11b^+^ DCs. Indeed, when we infected murine macrophages with *C. albicans* and measured fungal killing using a previously reported method [13], the *ole2/ole2* mutant exhibited reduced resistance to macrophages relative to the WT strain, and this defect in resistance to host killing can be reversed by the provision of exogenous PGE_2_ (Fig 5E-F).

**Fig 5.**
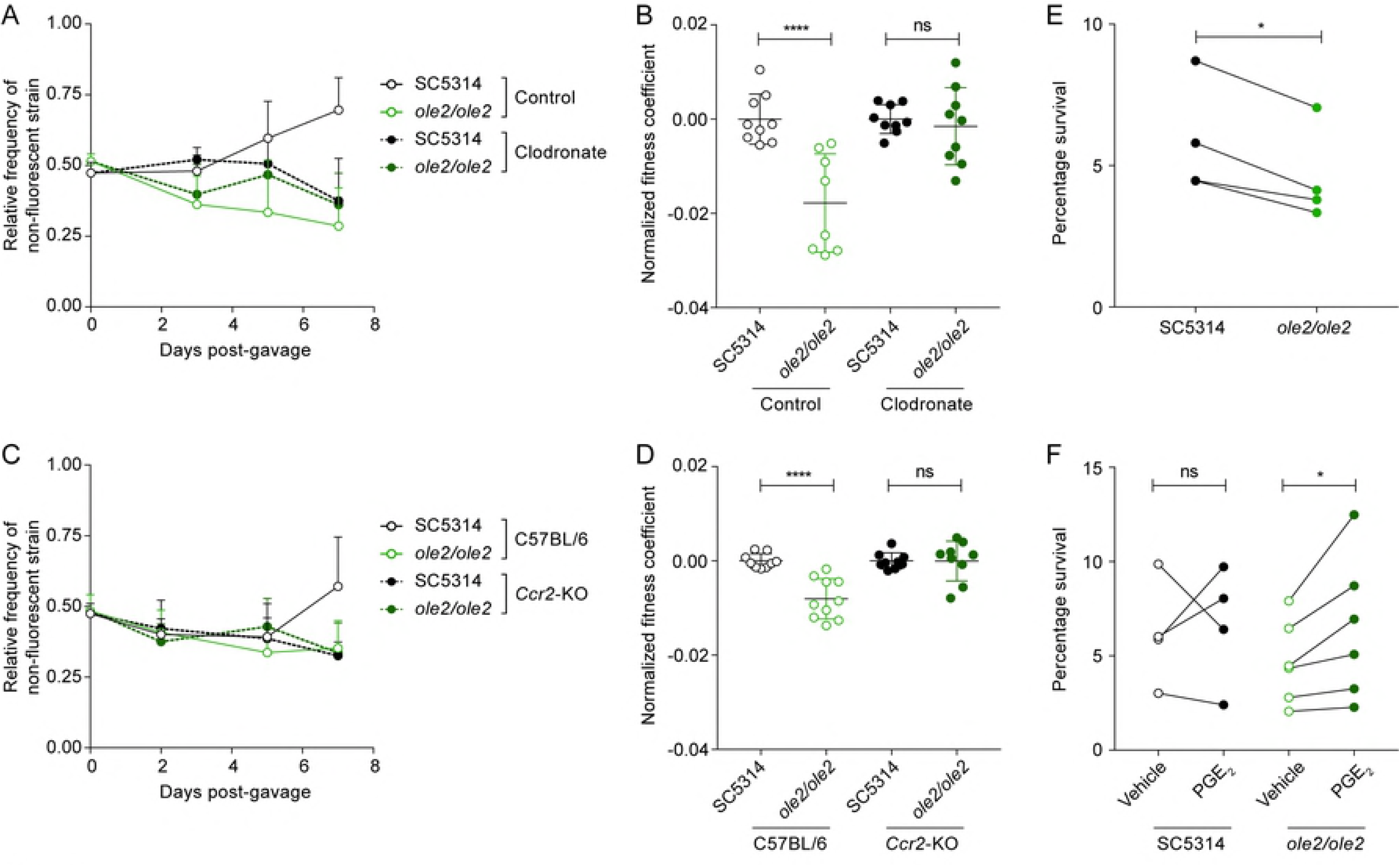
Ablation of colonic phagocytes abrogates the fitness defect of the *ole2/ole2* mutant. *In vivo* intestinal colonization experiments were performed as in Fig 4. (A-B) Mice were treated with control or clodronate-containing liposomes. (C-D) WT or *Ccr2*-KO mice were used for intestinal colonization. (A, C) Kinetics of the relative frequencies of the indicated fungal strains. (B, D) Fitness coefficients of the indicated fungal strains determined as described in Materials and Methods. Coefficients for each treatment group were normalized to the mean of the respective WT coefficients. (E) J774A.1 murine macrophages were infected with the indicated *C. albicans* strains for 24 h and the percentage survival of each strain was determined as outlined in Materials and Methods. (F) Infection of macrophages as in (A), but with addition of exogenous PGE_2_ or vehicle to the cultures. Data pooled from 2 (A-D) or 4-6 (E-F) independent experiments. Mean ± s.d. (A-D). *n*=9-10 mice/group (A-D). ****, *p* < 0.0001; ***, *p* < 0.001; *, *p* < 0.05; ns, not significant. Two-way ANOVA with Sidak’s multiple comparisons test (A-D); paired student’s *t* test (E-F).

## Discussion

In this study, we sought to clarify the physiological function of PGE_2_ production by *C. albicans*, a phenomenon long described [31] but of unclear importance to either host or fungus. Contrary to previous reports, using a PGE_2_-deficient fungal mutant, we found no contribution of fungal PGE_2_ to hyphal morphogenesis or fungi-induced macrophage cytotoxicity *in vitro*. Neither did fungal PGE_2_ influence the *in vivo* virulence of *C. albicans* across disparate murine models of candidiasis that affect different organs and engage distinct antifungal immunological responses. Instead, we uncovered a specific role for PGE_2_ production by *C. albicans* in potentiating fungal colonization of the murine intestine. Interestingly, the effects of fungal PGE_2_ are mediated by host phagocytes. Our results raise several questions that merit further elaboration.

How has *C. albicans* evolved the ability to synthesize PGE_2_ from AA, a substrate that it lacks? No fungal orthologs of the mammalian biosynthetic enzymes for PGE_2_ – the cyclooxygenases, COX-1 and COX-2, and the PGE_2_ synthase, PTGES – have been identified to date [36, 47]. Even using a sensitive method based on profile hidden Markov models (HMMER v.3) failed to identify distant homologs of the mammalian PGE_2_ biosynthetic enzymes (F. L. Sirota & S. Maurer-Stroh, unpublished observation). In addition, the three genes identified to contribute to PGE_2_ synthesis in *C. albicans* – *OLE2*, *FET3*, and *FET31* – do not harbor enzymatic domains that are known to bind to or act on AA or downstream prostaglandins. Indeed, *FET3* and *FET31* are putative iron transporters in *C. albicans* and might instead modulate fungal PGE_2_ production indirectly, possibly by acting as cofactors or regulating cellular levels of iron that in turn impact PGE_2_ synthesis. Genetic deficiency of *OLE2* in *C. albicans* did not completely eliminate PGE_2_ production *in vitro* [36] (Fig 1), implying the existence of additional components of the fungal PGE_2_ synthetic pathway. Thus *C. albicans* has convergently evolved a mechanism to generate PGE_2_ from a host-derived precursor, presumably in response to evolutionary pressures exerted by its lifestyle as an obligate symbiont in warm-blooded organisms.

Why does *C. albicans* goes through the effort of producing PGE_2_ at all? Earlier studies have implicated fungus-derived PGE_2_ in pathogenesis, since PGE_2_ promotes virulence traits such as germ tube [34] and biofilm [48] formation *in vitro*. Moreover, fungal PGE_2_ can suppress host immunity, which might in turn favor fungal growth. For instance, PGE_2_ emanating from a fungal bloom in the gut induced an immunoregulatory M2 phenotype in alveolar macrophages, which in turn exacerbated allergic airway inflammation in mice [32], and exposure to fungal PGE_2_ diminished the protective immune response generated in response to immunization with DCs pulsed with *C. albicans* yeasts [1, 2, 4]. However, the preceding findings do not point to an overt benefit for *C. albicans*, for which symbiosis is the *status quo* and infection is adventitious and often a byproduct of sustained immunodeficiency and/or microbial dysbiosis. Accordingly, we observed no role for fungal PGE_2_ in mediating the virulence of *C. albicans* under both *in vitro* and *in vivo* settings. Conversely, fungal PGE_2_ production appeared to be important for promoting intestinal colonization by *C. albicans*. *C. albicans* is located at high titers in the gut [6, 35] and acquisition of the fungus often occurs early during ontogeny via transmission of maternal intestinal strains to the infant [19, 20]. It would thus be evolutionarily advantageous for *C. albicans* to evolve an independent means of generating PGE_2_ to facilitate its own entrenchment as a symbiont in the gut.

How does fungal PGE_2_ mediate intestinal colonization by *C. albicans*? Here we provide evidence in support of gut phagocytes, in particular macrophages and CD11b^+^ DCs, as a target of fungal PGE_2_, since depletion of colonic phagocytes by two orthogonal approaches abrogated the fitness defect of the *ole2/ole2* mutant vis-à-vis the WT strain. The *ole2/ole2* mutant also sustained increased killing by macrophages *in vitro*, a phenomenon that was reversed by the addition of exogenous PGE_2_. PGE_2_-dependent fungal resistance to phagocytes could occur via the paracrine activity of fungal PGE_2_ on proximal phagocytes, and/or an autocrine effect of fungal PGE_2_ on the fungus itself. PGE_2_ has wide-ranging effects on phagocytes, most of which converge on immunosuppression [49]. For instance, PGE_2_ reduces the phagocytosis of fungi by macrophages [32, 50, 51] and polarizes both fungi-exposed macrophages and DCs towards an immunoregulatory phenotype, characterized by a decreased secretion of pro-inflammatory cytokines such as tumor necrosis factor (TNF)-α and an increased expression of suppressive factors such as interleukin (IL)-10 and arginase-1 [21–27]. Hence, PGE_2_ secreted by *C. albicans* proximal to or engulfed by intestinal phagocytes might dampen their uptake of more fungi, phagosome maturation, and/or production of inflammatory, antifungal factors. The effects of fungal PGE_2_ can also radiate beyond those of phagocytes to T cell responses primed by fungi-experienced phagocytes [52], which in turn control fungal growth in the gut. PGE_2_ is labile and likely has short-range activity, possibly even at the level of individual fungi-containing phagosomes in phagocytes [53, 54]. This might account for the apparent cell-autonomous effect of fungal PGE_2_ on intestinal fitness of *C. albicans*.

In addition, PGE_2_ might act directly on *C. albicans* itself to augment its resistance to host immunity. Mucosal microbial communities, including those of *Candida*, have been likened to biofilms [12], and certain genes crucial for the intestinal fitness of *C. albicans* are also required for biofilm formation [62], indicating common mechanisms underlying fungal fitness in both biofilms and the gut. Fungus-derived PGE_2_ might therefore elicit biofilm-like transcriptional programs in gut fungi that enhance fitness via various means, such as the induction of stress response pathways that improve fungal resistance to immunological assaults from the host [23, 24].

Our study focused on the role of fungus-derived PGE_2_, but does not rule out a role for host-derived PGE_2_ or other AA-derived lipid mediators in modulating intestinal symbiosis by *C. albicans*. Host phagocytes secrete PGE_2_ when stimulated with *C. albicans* [29, 63], though it is plausible that host- and fungus-derived PGE_2_ exert distinct activities in a tissue- and cell-type-specific fashion, especially in light of the differential accessibility of immunocyte populations to fungi and the instability of PGE_2_. Nevertheless, we predict that the effects of fungal PGE_2_ on the intestinal fitness of *C. albicans* would be amplified in the absence of host-derived PGE_2_. Notably, *C. albicans*, as well as other fungi and parasites, produces immunomodulatory lipid mediators from AA aside from PGE_2_ *in vitro*, including prostaglandin D2 and leukotrienes [26, 64, 65], some of which have been implicated in modulating microbial fitness during pathogenesis [7–14]. It is tempting to speculate that non-PGE_2_ lipids can, akin to PGE_2_, modulate the fitness of *C. albicans* and other symbiont microbes in the intestines and other barrier surfaces.

We report here a novel mechanism used by *C. albicans* to colonize the gut that is distinct from previously delineated pathways, such as the suppression of filamentation and changes in nutrient metabolism and iron acquisition [66]. Since fungal colonization is a prerequisite for disseminated or mucosal diseases, this work provides new potential targets for therapeutic control of intestinal fungal populations to prevent human fungal infections.

## Materials and Methods

### Ethics Statement

All animal experiments were approved by the BRC Institutional Animal Care and Use Committee (IACUC) in accordance with the guidelines of the Agri-Food & Veterinary Authority (AVA) and the National Advisory Committee for Laboratory Animal Research (NACLAR), Singapore, under IACUC protocol number 171271.

### Fungal strains

Fungal strains used in this study are listed in S1 Table. All fungal strains were grown in yeast peptone dextrose (YPD) media (1% (w/v) yeast extract, 2% (w/v) peptone, 2% (w/v) glucose) overnight at 30 °C prior to use in experiments unless otherwise stated.

### Construction of fungal strains

Fungal mutants were generated using the *SAT1*-flipping strategy as previously described [13, 67]. Briefly, the plasmid pSFS2 containing the *SAT1* flipper cassette was used to disrupt the open reading frame of the *OLE2*, *FET3*, and *FET31* genes of *C. albicans*. Sequences (500-600 bp) flanking the targeted genes were amplified using the primers listed in S2 Table. Gene deletion was verified by quantitative polymerase chain reaction (qPCR) using the primers listed in S2 Table.

### Mice and treatments

C57BL/6J and *Ccr2*-KO (B6.129S4-*Ccr2^tm1Ifc^*/J) mice were maintained under specific-pathogen-free conditions at the A*STAR Biological Resource Centre (BRC) in Biopolis, Singapore. Adult mice (6-8 weeks of age) of either sex were used in all experiments. To deplete phagocytes, mice were injected intraperitoneally (i.p.) with control or clodronate-containing liposomes (Liposoma B.V.) every 2 days starting from 2 days prior to the inoculation of mice with fungi. To perform PGE_2_ supplementation, mice were injected i.p. with 16.7 μg/kg of 16,16-dimethyl-PGE_2_ (Cayman Chemical) or vehicle (methyl acetate [Sigma]) daily, starting from day 0, throughout the course of fungal colonization.

### Measurement of fungal PGE_2_ production

Fungal PGE_2_ production was measured by enzyme-linked immunosorbent assay (ELISA) as previously described [68]. Briefly, fungi were inoculated into fresh YPD to an optical density (OD) at 600 nm of 0.2 and grown at 37 °C with shaking at 250 rpm for 2-2.5 h before peroxide-free arachidonic acid (Cayman Chemical) was added to a final concentration of 0.5 mM. Fungal growth was continued for another 48 h before supernatants were harvested. Levels of PGE_2_ metabolite (PGEM), a stable derivative of PGE_2_, in fungal supernatants were assayed using the PGEM EIA kit (Cayman Chemical) as per manufacturer instructions.

### Quantification of fungal titers

Fungal titers for all experiments were determined by plating fungal cultures, tissue homogenates, or lavage fluids on YPD agar containing 50 U/ml penicillin (Gibco) and 50 μg/ml streptomycin (Gibco). Plates were incubated at 37 °C for 24-48 h and colony-forming-units (CFUs) enumerated.

### Murine infection with *C. albicans*

To elicit systemic candidiasis, mice were injected intravenously via the tail vein with a sublethal dose of *C. albicans* (2 × 10^5^ cells). Oropharyngeal candidiasis (OPC) was established as previously described [69]. Briefly, mice were injected subcutaneously (s.c.) with 100 mg/kg cortisone 21-acetate (Sigma) in 0.05% (v/v) Tween®-80 (Sigma)/phosphate buffered saline (PBS) 1 day prior to infection to induce transient immunosuppression. Mice were subsequently sedated with 150 mg/kg ketamine and 10 mg/kg xylazine before sublingual inoculation with a small cotton ball saturated with *C. albicans* cells (2 × 10^7^ cells/ml) for 75 min. Vulvovaginal candidiasis (VVC) was established as previously described [68]. Briefly, female mice were treated s.c. with 0.1 mg of β-estradiol 17-valerate (Sigma) in sesame oil (Sigma) 3 days prior to infection. β-estradiol injections were maintained weekly thereafter. Mice were intravaginally inoculated with 5 × 10^4^ *C. albicans* cells in 20 μl of PBS. Infected mice were monitored daily (systemic candidiasis and OPC) for survival and weight loss. Tongues from mice with OPC were homogenized by pressing the tissues through a 40 μm cell strainer. Homogenates were plated to assess fungal burden. Mice with VVC were intravaginally lavaged with 100 μl of PBS. Vaginal lavage fluid was plated to determine fungal burden or used for neutrophil quantification by flow cytometry as described below under “Antibodies and flow cytometry”.

### Intestinal colonization with *C. albicans*

3-4 days prior to inoculation, mice were placed on drinking water containing 1 g/l of penicillin G (Sigma) and 2 g/l of streptomycin (Sigma) [13]. Antibiotics were changed twice weekly and maintained throughout the course of the experiment. Mice were gavaged with 10^7^ *C. albicans* cells to establish intestinal colonization. To determine the intestinal fungal load, stools were collected, weighed, homogenized in PBS, and plated.

### *In vivo* intestinal competition

Intestinal fungal competition experiments were performed as previously described [13]. Briefly, mice were gavaged with 2 × 10^7^ cells comprising the SC5314-dTomato strain and the test strain (WT or *ole2/ole2*) at a 1:1 ratio. The relative frequency of the test strain at each time point was determined by counting the number of fluorescent versus non-fluorescent colonies under a fluorescence stereomicroscope (Olympus). The fitness coefficient of each test strain in each mouse was calculated as previously described [13]. To control for inter-experimental variability, fitness coefficients were normalized to the mean of the coefficients for the WT strain within each treatment group (e.g. clodronate-treated group) for each experiment. Normalization was performed by calculating the difference between each fitness coefficient value and the mean for the WT strain.

### *In vitro* competition

*In vitro* competition experiments were performed as previously described [13]. Briefly, 1.25 × 10^6^ cells comprising the SC5314-dTomato strain and the test strain (WT or *ole2/ole2*) at a 1:1 ratio were inoculated into 50 ml of media (YPD or yeast nitrogen base [YNB] (BD Difco) with amino acids) and grown at 37 °C. Every 8 or 16 h, fungal cultures were harvested, plated, and 1.25 × 10^6^ cells re-inoculated into 50 ml of fresh media as described above. Relative proportions and fitness coefficients of each test strain were derived as described above.

### Infection of macrophages with *C. albicans*

Fungal resistance to macrophages was determined as previously described [13]. Briefly, confluent J774A.1 macrophages (ATCC) were infected with *C. albicans* cells at low multiplicities of infection (MOIs) ranging from 0.005 to 0.0003125 to ensure that essentially 100% of fungi were engulfed by the macrophages at 3 h post-infection. Infected macrophages were incubated at 37 °C and fungal colonies enumerated after 24 h. Fungi were also seeded in wells without macrophages at varying densities to assess the number of CFUs plated in each experiment. The percentage of fungal cells that escaped killing by macrophages was calculated as previously described [70]. Where indicated, 100 nM of PGE_2_ (Cayman Chemical) or the corresponding volume of vehicle (ethanol) was added to the cultures.

### Filamentation assays

Filamentation of *C. albicans* strains was assessed as previously described [13]. Briefly, *C. albicans* cells were incubated in Dulbecco’s Modified Eagle’s Medium (DMEM) supplemented with 10% fetal calf serum (FCS) at 37 °C in 96-well plates for 24 hours, or plated on Spider agar (1% nutrient broth, 1.35% agar, 0.4% potassium phosphate, 2% mannitol) at 37 °C for 4 days before colonies were scored for their morphology, i.e. wrinkled versus smooth, indicative of filamentous versus non-filamentous cells, respectively.

### *In vitro* cytotoxicity assay

Confluent J774A.1 macrophages were infected with *C. albicans* cells at a MOI of 1 for 6 h. LDH released into the culture media was measured using the CytoTox 96^®^ Non-Radioactive Cytotoxicity Assay (Promega) as per kit instructions and normalized to LDH levels in uninfected cells lysed with 1% Triton X-100 (Sigma).

### Measurement of AMP production from IECs

Confluent human colonic epithelial cell lines HT-29 and Caco-2 (both from ATCC) were infected with *C. albicans* cells at a MOI of 0.5 for 24 h. LL-37 secreted into the culture media was measured by the LL-37 ELISA kit (Cusabio) as per manufacturer instructions.

### Assessment of fungal resistance to mCRAMP

Fungal resistance to mCRAMP was evaluated as previously described [13]. Briefly, overnight cultures of *C. albicans* were inoculated into fresh YPD in the presence of 400 nM PGE_2_ (Cayman Chemical) or vehicle (ethanol) for 3 h. Cells were washed, resuspended in the appropriate concentration of mCRAMP_1-39_ (Synpeptide), and incubated at 37 °C for 2 h. Cells were washed extensively after mCRAMP exposure and stained with 50 μM of propidium iodide (PI) (Thermo Fisher Scientific) for 15 min. The proportion of live PI^−^ fungal cells was determined by flow cytometry.

### Assessment of fungal β-glucan exposure

Exposure of β-glucan on fungal cell walls was assessed by flow cytometry and a hDectin-1b reporter cell line as previously described [13]. Briefly, overnight cultures of *C. albicans* were inoculated into fresh YPD for 4 h. Cells were harvested and fixed with 1% neutral buffered formalin (Sigma) for 20 min and washed extensively. A portion of the fungal cells was blocked with 1% bovine serum albumin (BSA) (Capricorn Scientific) prior to staining for β-glucan as outlined under “Antibodies and flow cytometry”. A second portion of cells was co-cultured with the HEK-Blue™ hDectin-1b reporter cell line (InvivoGen), which secretes alkaline phosphatase (SEAP) in amounts commensurate with the strength of hDectin-1b-NF-κB signaling, at a 10:1 ratio (fungi:reporter cell). A hDectin-1b^−^ control cell line, HEK-Blue™ Null2 (InvivoGen), was concomitantly stimulated with fungi. After incubation at 37 °C for 18-24 h, SEAP levels in the culture supernatant were quantified using QUANTI-Blue™ (Invivogen) as per manufacturer instructions. SEAP activity in supernatants derived from the hDectin-1b cell line was normalized to that from the Null2 line.

### Measurement of transcript levels by quantitative PCR

RNA was extracted from a small segment of the distal colon harvested from *C. albicans*-colonized mice 7 days after inoculation. Samples were homogenized in TRIzol™ (Invitrogen) using 0.5 mm glass beads (Sigma) and RNA extracted as per manufacturer instructions. cDNA was prepared from RNA using SuperScript™ III reverse transcriptase (Invitrogen) as per manufacturer instructions. Levels of *Camp* in cDNA were measured by qPCR using PerfeCTa SYBR^®^ Green FastMix (Quantabio) and normalized to those of *Rpl13a*.

### Isolation of colonic lamina propria cells

To remove the epithelial layer, colons were sliced open longitudinally to expose the luminal surface and shaken in DMEM containing 1 mM dithiothreitol, 5 mM EDTA, and 3% (v/v) FCS for 20 min at 37 °C. Remaining tissue was washed in cold DMEM, minced, and digested in DMEM containing 1.5 mg/ml collagenase D (Roche), 50 μg/ml DNase I (Roche), and 2% (v/v) FCS for 45 min at 37 °C. Digested tissue was washed and filtered twice to obtain a single-cell suspension.

### Antibodies and flow cytometry

Mammalian cells were stained with antibodies against CD3ε (145-2C11), CD4 (RM4-5), CD8α (53-6.7), CD11b (M1/70), CD11c (N418), CD19 (6D5), CD45 (30F11), CD64 (X54-5/7.1), I-A/E (M5/114.15.2), Ly6G (1A8) from Biolegend and 4’,6-diamidino-2-phenylindole (DAPI) (Thermo Fisher Scientific) to exclude dead cells. All immunocyte subsets were gated on live leukocytes (DAPI^−^CD45^+^). Macrophages, DCs, B cells, CD4^+^ T cells, and CD8^+^ T cells from the colonic lamina propria were defined as CD64^+^CD11b^+^CD11c^+^I-A/E^+^, CD64^−^CD11c^+^I-A/E^+^, CD3ε^−^CD19^+^, CD3ε^+^CD19^−^CD4^+^, and CD3ε^+^CD19^−^CD8α^+^, respectively. Intravaginal neutrophils were defined as CD11b^+^Ly6G^+^ cells. Fungal cells were stained with anti-β-1,3-glucan (Biosupplies) followed by Alexa Fluor^®^ 647-conjugated goat anti-mouse IgG (Invitrogen). Samples were acquired on the MACSQuant^®^ VYB (Miltenyi Biotec) and analysed using FlowJo v10.2 (Tree Star).

### Statistics

Pairwise comparisons were performed using the Student’s *t* test or the paired Student’s *t* test, as appropriate, while multi-group comparisons were performed using the Two-way Analysis of Variance (ANOVA) with Sidak’s multiple comparisons test (Prism 7; Graph-Pad), unless otherwise stated. *P* values were deemed significant if less than 0.05. Data routinely presented as mean ± standard deviation (s.d.) unless otherwise stated.

## Acknowledgments

We thank G.T.T. Le for technical assistance; S. Maurer-Stroh and F. L. Sirota for sharing unpublished bioinformatics analyses; G. Rancati for scientific discussions; Singapore Immunology Network (SIgN) Mouse Core for providing mice. This paper is dedicated to the memory of Royston Leong.

## Supporting information

**S1 Fig. Parameters of *in vivo* virulence of *C. albicans*.**

(A-B) Weight change of mice infected systemically (A) or sublingually (B) with the indicated *C. albicans* strains. (C) Neutrophil counts in vaginal lavage fluid over the course of vulvovaginal infection with the indicated fungal strains. Data pooled from 2 independent experiments. Mean ± s.d.. *n*=8-10 mice/group. Student’s *t* test.

**S2 Fig. Fungal PGE_2_ promotes intestinal colonization by *C. albicans*.**

Mice were inoculated with WT or the *ole2/ole2* mutant of *C. albicans* in competition with a fluorescently-tagged WT strain as described in Fig 4. Mice were left untreated (A) or supplemented daily with 16,16-dimethyl-PGE_2_ (16,16-dmPGE_2_) or its corresponding vehicle (B-C). Fitness coefficients (A, C) and relative frequencies (B) of the indicated fungal strains across independent experiments. Mean ± s.d.. *n*=4-5 mice/group per experiment. ***, *p* < 0.001; **, *p* < 0.01; *, *p* < 0.05; Student’s *t* test.

**S3 Fig. Fungal PGE_2_ does not alter host AMP production or fungal resistance to AMPs.**

(A) The human IEC cell lines HT-29 and Caco-2 were infected with the indicated *C. albicans* strains at a MOI of 0.5 for 24 h. Secretion of LL-37 was measured by ELISA and normalized to that of uninfected cells (media only). (B) Transcript levels of *Camp* in the colonic tissue of mice colonized with the indicated fungal strains. (C) The viability of the indicated fungal strains after exposure to mCRAMP as assessed by propidium iodide staining. Fungal strains were grown in exponential phase in the presence or absence of PGE_2_ prior to incubation with mCRAMP. Data pooled from 2 independent experiments. *n*=6-8 (B). Mean ± s.d.. ns, not significant. Student’s *t* test.

**S4 Fig. Effects of fungal PGE_2_ on β-glucan exposure and dectin-1 signaling.**

Exponentially growing *C. albicans* strains were stained for (A) surface β-1,3-glucan, or (B) used to stimulate HEK-Blue™ hDectin-1b reporter cells, which transduce hDectin-1b signaling into NF-κB-driven alkaline phosphatase secretion. hDectin-1b signaling strength is quantified by measuring the activity of alkaline phosphatase secreted by the reporter cells. (A) The mean fluorescence of surface β-glucan on the indicated fungal strains was normalized to that of the WT strain. (B) The strength of hDectin-1b signaling triggered by the indicated fungal strains, normalized to that of the WT strain. Data pooled from 6-8 independent experiments. Mean ± s.d.. *, *p* < 0.05; ns, not significant. Paired *t* test.

**S5 Fig. Numbers of various immunocyte populations in the colonic lamina propria of mice depleted of phagocytes 7 days after *C. albicans* gavage.**

(A-B) Mice treated with control or clodronate-containing liposomes. (C-D) WT versus *Ccr2*-KO mice. MΦ, macrophage. Data pooled from 2 independent experiments. Mean ± s.d.. *n*=4-10 mice/group. ****, *p* < 0.0001; **, *p* < 0.01; *, *p* < 0.05. Student’s *t* test.

**S6 Fig. Ablation of colonic phagocytes abrogates the fitness defect of the *ole2/ole2* mutant.**

*In vivo* intestinal colonization experiments were performed as described in Fig 5 and fungal fitness coefficients for each experiment were determined as outlined in Materials and Methods. (A-B) Mice were treated with control or clodronate-containing liposomes. (C-D) WT or *Ccr2*-KO mice were used for intestinal colonization. Data pooled from 2 independent experiments. Mean ± s.d.. *n*=4-5 mice/group per experiment. ****, *p* < 0.0001; **, *p* < 0.01; *, *p* < 0.05; *#, p* = 0.059; Two-way ANOVA with Sidak’s multiple comparisons test.

**S1 Table.**

Fungal strains used in this study.

**S2 Table.**

Primers used in this study.

